# European-wide forest monitoring substantiate the neccessity for a joint conservation strategy to rescue European ash species (*Fraxinus spp*.)

**DOI:** 10.1101/2021.07.28.454255

**Authors:** Jan-Peter George, Tanja GM Sanders, Volkmar Timmermann, Nenad Potočić, Mait Lang

## Abstract

European ash (*Fraxinus excelsior*) and narrow-leafed ash (*F. angustifolia*) are keystone forest tree species in Europe with a broad ecological amplitude and significant economic importance. Besides global warming both species are currently under significant thread by an invasive fungal pathogen that has been progressively spreading throughout the continent for almost three decades. Ash dieback caused by the invasive ascomycete *Hymenoscyphus fraxineus* is capable of damaging ash trees of all age classes and often leads to the ultimate death of a tree after years of progressively developing crown defoliation. While studies at national and regional level already suggested rapid decline of ash populations as a result of ash dieback, a comprehensive survey at European level with harmonized crown assessment data across countries could shed more light into the population decline from a pan-European perspective and could also pave the way for a new conservation strategy beyond national boarders. Here, we present data from the ICP Forests Level I crown condition monitoring including 27 countries, covering the timespan from 1987-2020. In total, 407 survey plots randomly distributed across these countries were analyzed resulting in >36,000 individual observations. We found a substantial increase in defoliation and mortality over time indicating that crown defoliation has almost doubled during the last three decades. Hotspots of mortality are currently situated in southern Scandinavia and north-eastern Europe, well corresponding to the fact that the disease spread fast from north-east to north-west. Overall survival probability after nearly 30 years of infection has already reached a critical value of 0.51, but with large differences among regions (0.00-0.907). Both a Cox proportional hazard model as well as an Aalen additive regression model strongly suggest that survival of ash is significantly lower in locations with excessive water regime and which experienced more extreme precipitation events during the last two decades. Our results underpin the neccessity for fast governmental acting and joint rescue efforts beyond national boarders since overall mean defoliation will likely reach 50% as early as 2030 as suggested by time series forecasting. We strongly recommend to develop a pan-European conservation strategy before the decline will reach its tipping point resulting into non-reversible loss of diversity in the European forest landscape.

## 1. Introduction

European ash (*Fraxinus excelsior*) and narrow-leafed ash (*F. angustifolia*) are keystone forest tree species with high ecological and economical value for the European forest sector (Hill et al. 2019). As light demanding pioneer species, they succesfully inhabit a wide range of different ecosystems including riparian forests and mountain forests amongst others (Pliura & Heuertz, 2003). Their ability to particularly tolerate temporary flooding makes them both indispensable components of alluvial forests with strongly fluctuating hydrological regimes (Dufour and Piegay, 2008). Moreover, both ash species are host to a wide range of other taxa including mammals, birds, invertebrates, and bryophytes. As such, Mitchell et al. (2014) estimated that from 953 species associated with European ash, 69 can be characterized as being “highly associated” with its host, which means that these species will be under threat of becoming rapidly extinct when the host population will decline.

However, European ash and narrow-leafed ash are currently endangered, since the invasive pathogen *Hymenoscyphus fraxineus* has entered Europe around 1990 (Przybył 2002). The disease is commonly known as ash dieback (hereafter abbreviated with „ADB”) and has devastatingly spread over entire Europe throughout the last 30 years (Vasaitis & Enderle, 2017) with an expansion velocity of 30-70 km per year (Børja et al. 2017; Queloz et al. 2017; Ghelardini et al. 2017; Marçais et al. 2016). The epicentre of the disease is thought to be located in north-eastern Poland and has concentrically expanded through wind-dispersed ascospores as well as by human activity (Orton et al. 2018). Briefly, leaves of ash trees are infected by windborne ascospores from where the fungus spreads via the petiole-shoot junction into the woody tissue of the tree. Recurrent infections cause rapid crown decline which often leads to mortality of trees after a few years (Enderle et al. 2019). Both species are thought to be equally susceptible against ADB (Enderle et al. 2019), while *Fraxinus ornus* as the third ash species occurring in Europe seems to be more resilient against the disease (Heinze et al. 2017) and is hence not considered in this study.

The extent to which the European population of *F. excelsior* and *F. angustifolia* has been declining since the arrival of the pathogen is largely unknown, which makes targeted conservation efforts at pan-European level challenging. National and regional reports on mortality rates were so far based upon literature reviews (e.g. Coker et al. 2019), experimental forest sites (Stocks et al. 2017, Cleary et al. 2017), and national forest inventory data (e.g. Díaz-Yáñez et al. 2020; Klesse et al. 2021; Enderle et al. 2018). However, the different approaches, datasets, and their specific sampling biases make it difficult to draw a conclusive pattern for European ash and narrow-leafed ash across their entire distributions. In particular, mapping mortality rates in space and time for entire Europe across the last three decades based on a consistent dataset could inform conservationists and policy makers at national and international level about the current status of ash and can unravel mortality hotspots which should be given highest priority for rescue and conservation programs in the near future. In this study we make use of survey data from the ICP Forests Level I network, providing detailed information on crown condition and diseases in annual resolution since 1987 (e.g. Timmermann et al. 2021). In total, we analyzed 36,170 observations on ash trees from 1987-2020 in 27 countries and incorporated stand information as well as climate data in order to test for covariation between ash dieback-induced mortality and ecological variables that may accelerate or decelerate the decline as was recently suggested in earlier studies at regional scale (e.g. Klesse et al. 2021; Díaz-Yáñez et al. 2020; Chumanova et al. 2019). We hypothesize that the mortality of both ash species has significantly increased throughout the last three decades resulting in a devastating ash dieback driven by ADB rather than by other abiotic and biotic agents. Moreover, we hypothesize that moist growing conditions, high abundance of ash, and moist weather conditions exacerbates mortality as they are likely to provide optimal growing conditions for the pathogen. Finally, we aim to use the past defoliation history to forecast the near future and unravel critical points for the long-term vitality and persistence for both species.

## 2. Material & Methods

### 2.1 Data

We used the ICP Forests Level I forest damage survey since this dataset constitutes the only available systematic survey sample across Europe at a 16×16km grid (Eichhorn et al. 2016). Within this dataset, crown parameters and damaging agents are assessed at an annual scale by survey teams typically including 24 dominant trees per plot. Crown defoliation is assessed in 5%-steps with 99% indicating complete defoliation of a tree and 100% its death. Damaging agents are assessed and are divided into abiotic, biotic and other damage causes. We selected all plots where at least one ash tree (either *F. excelsior* or *F. angustifolia*) was recorded and analysed the complete survey period spanning the years 1987 to 2020. An ash tree was subseuqently classified as dead when its defoliation status reached 100% and the tree did not occurr any longer in subsequent survey years.

#### 2.2.1 Overall survival probability after 30 years

Mortality data in tree ecology is often treated as a binary variable that can be best accomodated by logistic regression analysis (e.g. Koontz et al. 2021, Taccoen et al. 2019, Coker et al. 2019). However, when the cause of death is primarily caused by a pathogen which resembles a mortality pattern close to a pandemic situation (i.e. a time-to-death approach), logistic regression will be rather unsuitable. The reason being that such data is usually censored, which either means that at a specific survey date not all subjects had yet experienced the event (i.e. death) or got lost without any further assessments conducted (Therneau, 2020). Consequently, we used a different approach that specifically treats time-to-event data and that is usually applied in medical data such as survival analysis for patients carrying a lethal disease (e.g. Godaert et al., 2018; Sageran et al. 2008). We justify this because of two reasons: i) ash dieback usually constitutes a dead end for infected trees, since the disease is highly lethal and progressive and recovery of trees is rarely observed (Enderle et al. 2019), ii) ash dieback spreads through windborne ascospores at a fast rate so that we can assume that a large part of the ash population in Europe has already experienced high and equal infection pressure (Gross et al. 2014).

We first estimated the overall survival probability of ash with the Kaplan-Maier step function written as:

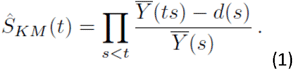

with S_KM_ being the overall survival probability, 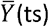 being the trees at risk at time t, d(s) being trees that died since start of infection and 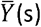 being the trees at risk since the beginning of infection. Since the observation plots are usually re-visited each year, the Kaplan-Maier curve will display the overall survival probability and 95% confidence intervals at one-year steps. We calculated the overall survival probability for two different models: i) Model 1 with simultanous infection: in the first model we assumed that the start of infection happened in 1992 for the entire European population. This is the first date when ash dieback was observed in northern Poland and it is generally believed that this is the epicentre of the disease (Enderle et al. 2019). ii) Model 2 with non-simultanous infection: Although the disease spread rapidly towards the west, east, and south from the epicentre, there were distinct time gaps between subsequent infections of ash populations located further apart from the epicentre. Hence, we gathered dates of first observations of ash dieback (Tab. 1) within the single countries from different literature resources (Vasaitis & Enderle, 2017; Solheim & Hietala, 2017). Even though these dates are to some extent biased and may do not reflect the exact date of arrival of ash dieback at national level, they can nevertheless serve as a good proxy for the temporal delay in infection due to expansion history of the disease. In few cases, the time between last assessment and event (nominator of equation 1) was negative in the second model, since some mortality already occurred before the estimated arrival of ash dieback. This affected only Spain and southern France, where some cases of mortality were atributable to other disturbance agents rather than to ash dieback (see discussion below). These few cases were consequently removed from both models.

**Table 1:**
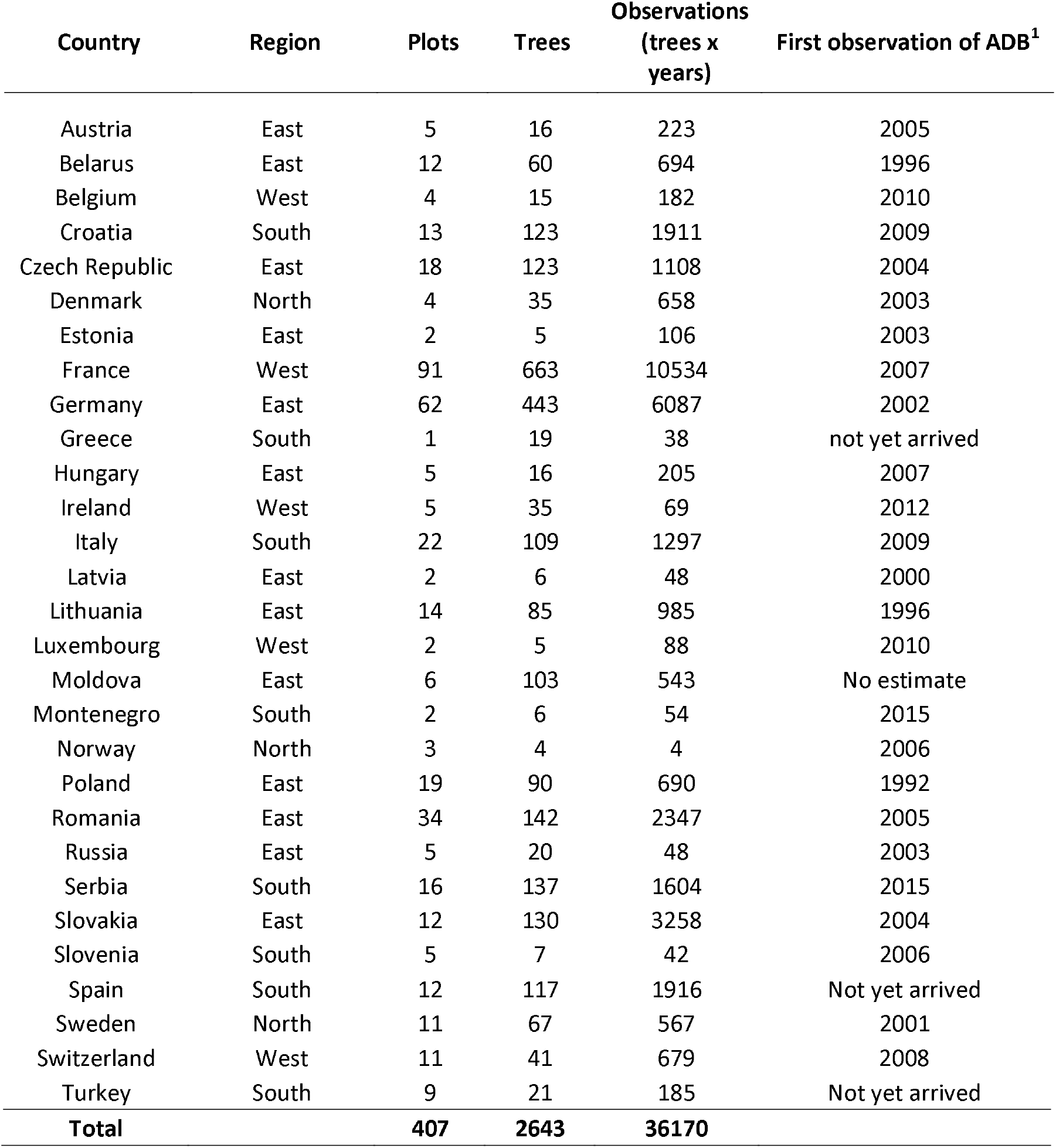
Summary of observations by country

| Country | Region | Plots | Trees | Observations<br>(trees x<br>years) | First observation of ADB<sup>1</sup> |
| --- | --- | --- | --- | --- | --- |
| Austria | East | 5 | 16 | 223 | 2005 |
| Belarus | East | 12 | 60 | 694 | 1996 |
| Belgium | West | 4 | 15 | 182 | 2010 |
| Croatia | South | 13 | 123 | 1911 | 2009 |
| Czech Republic | East | 18 | 123 | 1108 | 2004 |
| Denmark | North | 4 | 35 | 658 | 2003 |
| Estonia | East | 2 | 5 | 106 | 2003 |
| France | West | 91 | 663 | 10534 | 2007 |
| Germany | East | 62 | 443 | 6087 | 2002 |
| Greece | South | 1 | 19 | 38 | not yet arrived |
| Hungary | East | 5 | 16 | 205 | 2007 |
| Ireland | West | 5 | 35 | 69 | 2012 |
| Italy | South | 22 | 109 | 1297 | 2009 |
| Latvia | East | 2 | 6 | 48 | 2000 |
| Lithuania | East | 14 | 85 | 985 | 1996 |
| Luxembourg | West | 2 | 5 | 88 | 2010 |
| Moldova | East | 6 | 103 | 543 | No estimate |
| Montenegro | South | 2 | 6 | 54 | 2015 |
| Norway | North | 3 | 4 | 4 | 2006 |
| Poland | East | 19 | 90 | 690 | 1992 |
| Romania | East | 34 | 142 | 2347 | 2005 |
| Russia | East | 5 | 20 | 48 | 2003 |
| Serbia | South | 16 | 137 | 1604 | 2015 |
| Slovakia | East | 12 | 130 | 3258 | 2004 |
| Slovenia | South | 5 | 7 | 42 | 2006 |
| Spain | South | 12 | 117 | 1916 | Not yet arrived |
| Sweden | North | 11 | 67 | 567 | 2001 |
| Switzerland | West | 11 | 41 | 679 | 2008 |
| Turkey | South | 9 | 21 | 185 | Not yet arrived |
| **Total** |  | **407** | **2643** | **36170** |  |

#### 2.2.2 Cox proportional hazard model and Aalen additive regression model

In order to model the transition from „alive“ to „dead“ we used a Cox-proportional hazard model (Cox, 1972) and, alternatively, an Aalen additive regression model (Aalen, 1989). Such models are commonly applied in clinical testing for explaining mortality patterns for patients under risk when the data is right- or left-censored (Therneau & Grambsch, 2000). Both models are capable of incorporating covariates of different structure and specifically relate those covariates to survival patterns or any other censored outcome. The Cox-model has the form:

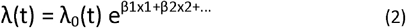

where λ(t) is the single-tree hazard as a function of time *t,* λ_0_ is the so-called baseline hazard, *X_1,2_* are the covariate vectors of subject *i* and β_1,2_ specify vectors of coefficients associated with the covariates (Therneau & Grambsch, 2000). The Aalen additive regression model was suggested as an alternative to the Cox-proportional hazard model, because some of the assumptions behind the Cox-model may not always be met by all included covariates (e.g. that all coefficients are constant over time) and because additive effects may reflect the nature of the underlying disease better than the proportionality assumption, in particular when the baseline hazard is very small. The additive model has the linear form:

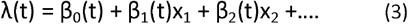

with β_0_ being the baseline function and β_i_ (t)x_i_ being the regression functions which determine the influence of the covariates. In contrast to the Cox-proportional hazard model, regression functions in (3) can freely vary over time.

We used two key stand characteristics from the ICP Forests Level I dataset (water availability coded as 1=insufficient, 2= sufficient, 3=excessive and humus type comprising 9 classes) as well as abundance of ash trees in each plot (number of ash trees/total number of trees) as survival predictors. The rationale behind the latter variable is that we hypothesize higher mortality rates when the host abundance is high compared to stands where only few ash trees occur.

Various recent studies showed that tree mortality, in particular in the very last years, has accelerated mainly as a result of climatically-driven changes such as warming and drought (e.g. Neumann et al. 2017; Senf et al. 2020). In order to disentangle whether the mortality pattern in ash between 1987 and 2020 has been driven stronger by ADB, climate or both, we calculated the monthly anomaly in maximum temperature and precipitation sums from 1995-2020 for all survey plots in which ash occurred. We used E-OBS daily gridded meteorological data at 0.25° spatial resolution (Haylock et al. 2008) and calculated the number of extreme events between 1995 and 2020 for each survey plot. Extreme events were defined as months which showed a standardized climatic anomaly of > or < two standard deviations above or below long-term average, respectively.

Since we expect that some covariates are not randomly distributed in space across the observation plots (i.e. the disease is generally stronger in the east compared to the west of Europe due to the infection history), we divided the data also into geographic strata so that each group (eastern, western, southern, and northern Europe) would have its distinct baseline hazard function, but common values for β. We employed the *survival* package in R (R Development Core Team, 2017) for all analysis steps (overall survival probability, Cox model, Aalens additive regression model) described above.

### 2.3 Time-series forecasting of crown defoliation

Finally, we aim to project the past defoliation history (1987-2020) of ash in Europe into the near future in order to determine critical time horizons for conservation and rescue management. We first calculated average defoliation per year and plot including all ash trees and performed timeseries forecasting spanning 50 years from the last available observation (2021-2070). We used both an ARIMA (autoregressive integrated moving average) model as well as exponential smoothing (Holts linear trend method according to Holt (1957)) as alternative approach. Briefly, a non-seasonal ARIMA model was fittted by determining the parameters p, d, and q where p is the order of the autoregressive part, d is the degree of first differencing, and q is the order of the moving average part. We first determined the three parameters by using the *auto.arima* function on the log transformed defoliation time series with the help of the *forecast* package in R (Hyndman & Khandakar, 2008). In order to evaluate whether the determined parameters were correctly chosen, we visually inspected the partial autocorrelation plot and checked the model residuals by performing a Portmanteau-test. We calculated 95% confidence intervals around both estimators (ARIMA, exponential smoothing) and evaluated the goodness of the obtained forecasts by using the first 22 years (1987-2009) as training dataset and the last ten years of observations (2010-2020) as evaluation data set. The mean square error (MSE) was used to compare forecast results between the two methods.

## 3. Results

### 3.1 Observed mortality and defoliation across Europe between 1987 and 2020

We identified 407 plots across 27 countries in which European ash or narrow-leafed ash occurred since the start of the survey in 1987 (Fig. 1, Tab. 1). This resulted in a total of 36,170 observations (plots x trees x years). Between 1987 and 2000 mortality frequency was moderate and occurred only sporadically in Spain, France, Romania, Slovakia, Italy, Lithuania and Moldova (range 0.006-0.15). Nevertheless, one stand in Belarus already showed 100% mortality between 1987 and 2000 (Fig. 2). Between 2000 and 2010 mortality accelerated mainly in Poland, Lithuania, Belarus, Sweden, Denmark, Germany, Czech Republic and at some local spots in France. While in most of these countries mortality rates were still low (1 to 3%), Poland and Sweden already showed significant mortality within stands (10% to 100%). However, in the following decade (2010-2020) ash mortality significantly accelerated with observed mortality in nearly all parts of Europe. Most notably, southern Scandinavia became a mortality hotspot with eight stands where all ash trees died within this period.

**Fig. 1:**
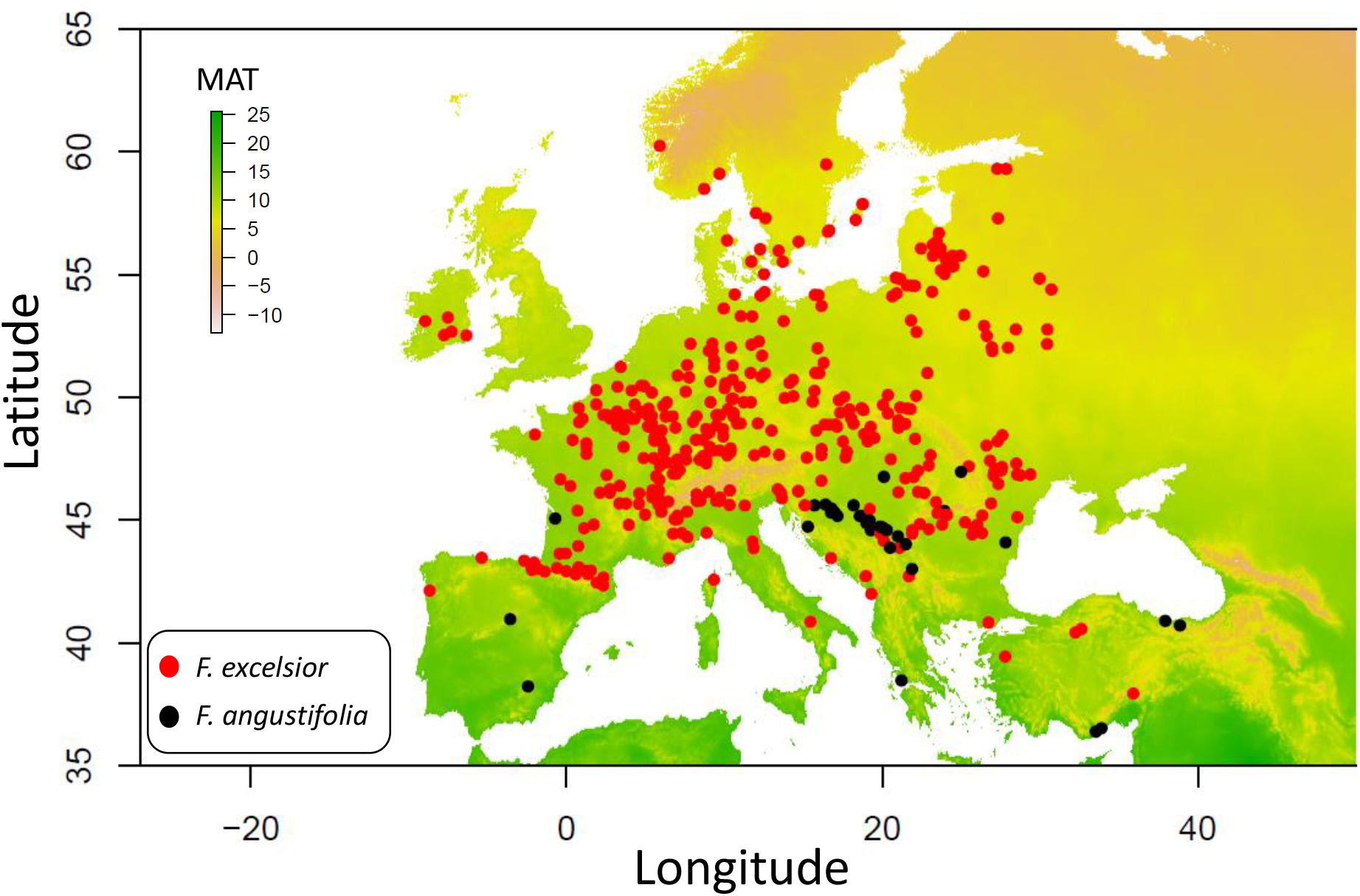
Overview of analzysed plots from he ICP Forests Level I dataset.

**Fig. 2:**
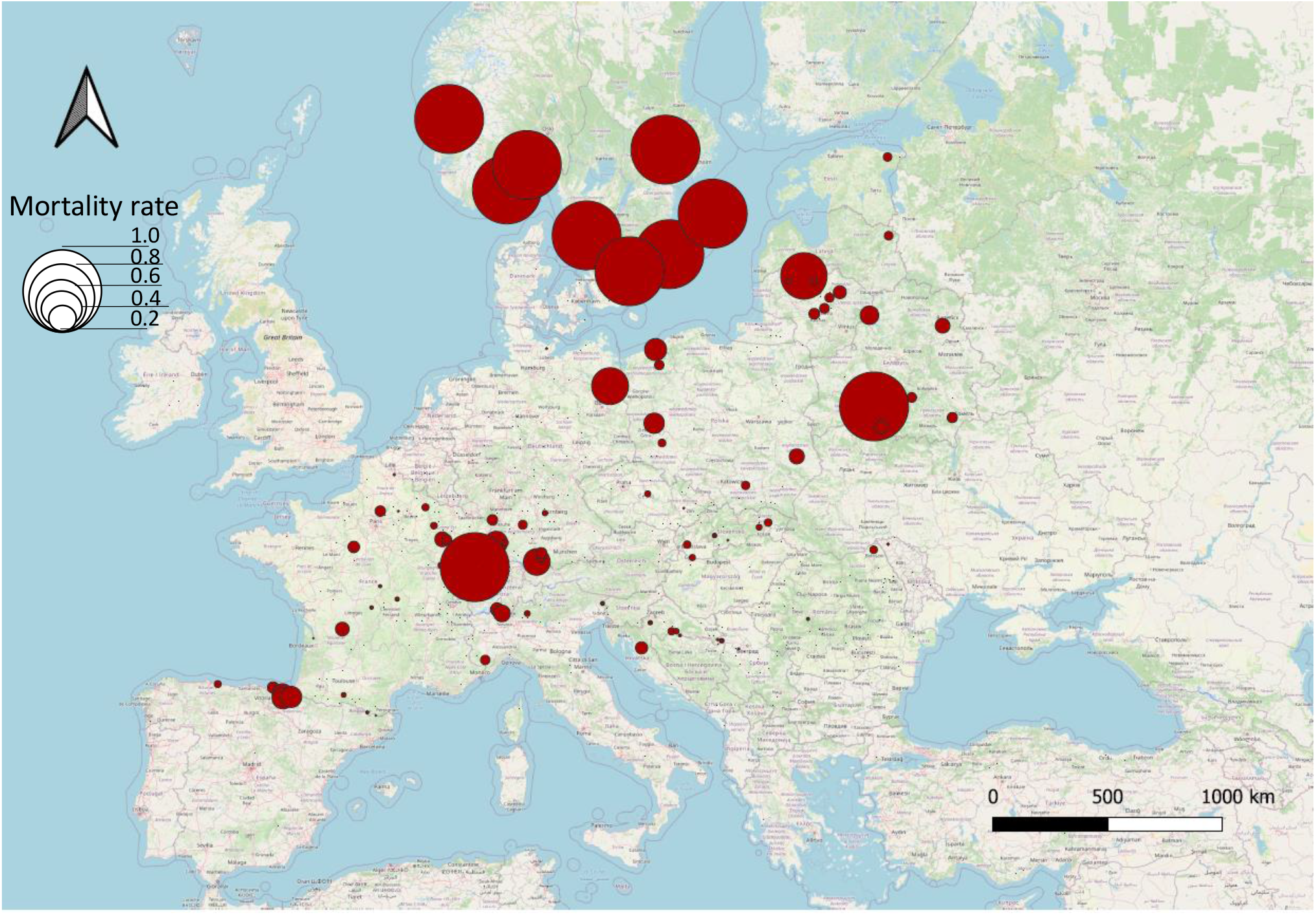
Cummulative mortality rate of ash between 1987 and 2020.

Generally, crown conditions of ash trees became progressively worse since the beginning of the survey across the entire range of occurrence. While average defoliation was 15% until the year 2000, it already reached 25% in 2010. After the latest survey in 2020 mean defoliation of European ash and narow-leafed ash has reached 38% on average. Mean defoliation showed strongest increase in the eastern, northern, and central part of the distribution since 1987 (Belarus, Denmark, Estonia, Latvia, Lithuania, Poland, Sweden, Germany, France, Luxembourg, Slovakia), but mostly non-significant trends in the southern part (Croatia, Moldova, Romania, Serbia, Slovenia, Spain, Switzerland, Turkey) (Fig. 3).

**Fig. 3:**
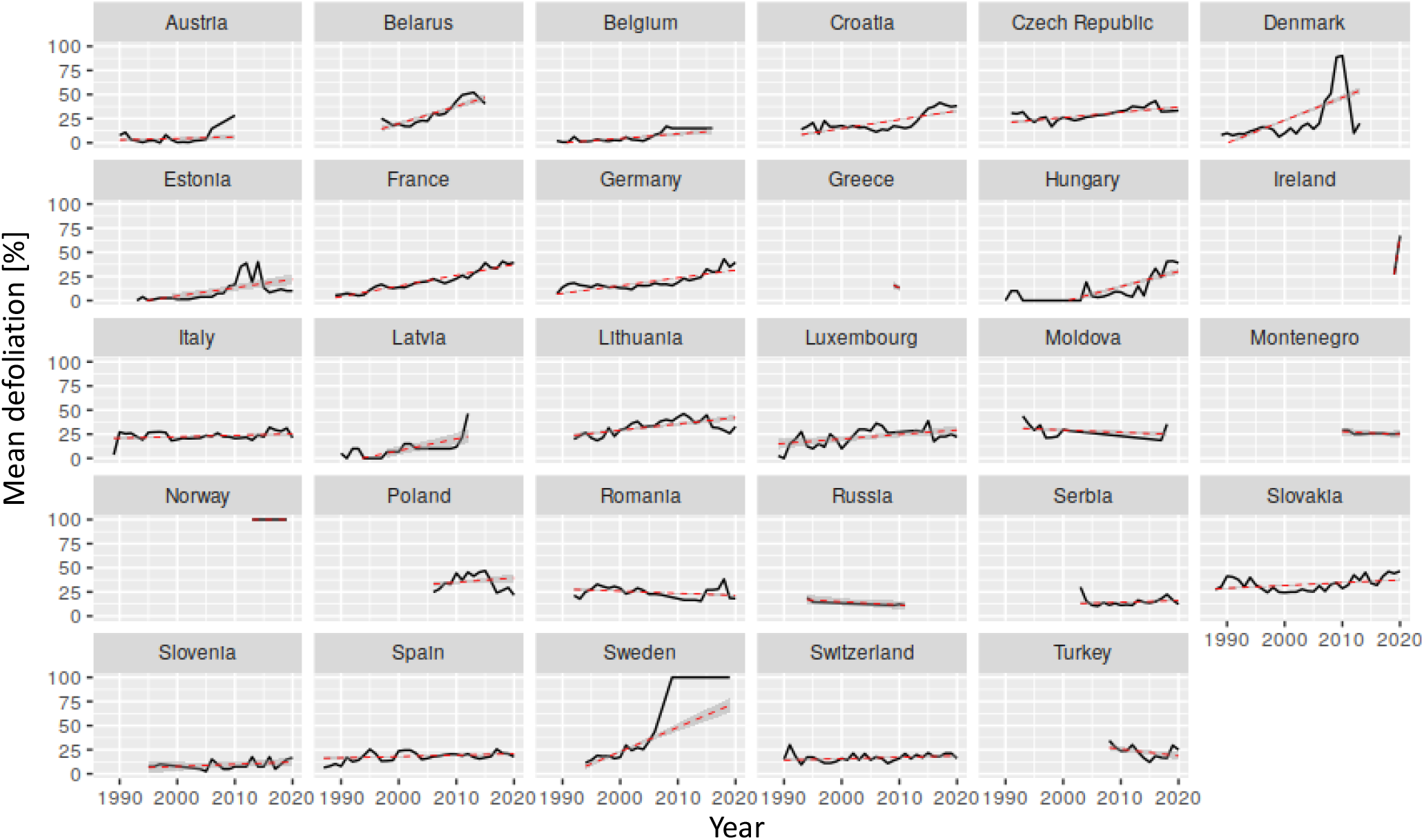
Mean defoliation across plots by survey country. Red dashed line shows linear time-series trend. Note that the downward trend in the Danish plots is caused by lost follow-up of some plots rather than recovery.

### 3.2 Survival probability after 30 years of infection history

The two different survival models (simultanous/non-simultanous infection) gave similar results for the first 10 years after infection, but diverged right after that period. After 20 years of infection model 1 which assumed an equal start of infection for all ash populations estimated a much higher survival probability of 0.95 (95%CI: 0.939-0.959) while model 2 (assuming that infection happened delayed) estimated an overall survival probability of 0.67 (0.607-0.734). Overall survival probability after the last assessment of trees in 2020 was 0.83 (0.805-0.846) for model 1 and 0.5 for model 2 (0.404-0.616). The Kaplan-Maier step functions for both models are shown in Fig. 4.

**Fig. 4:**
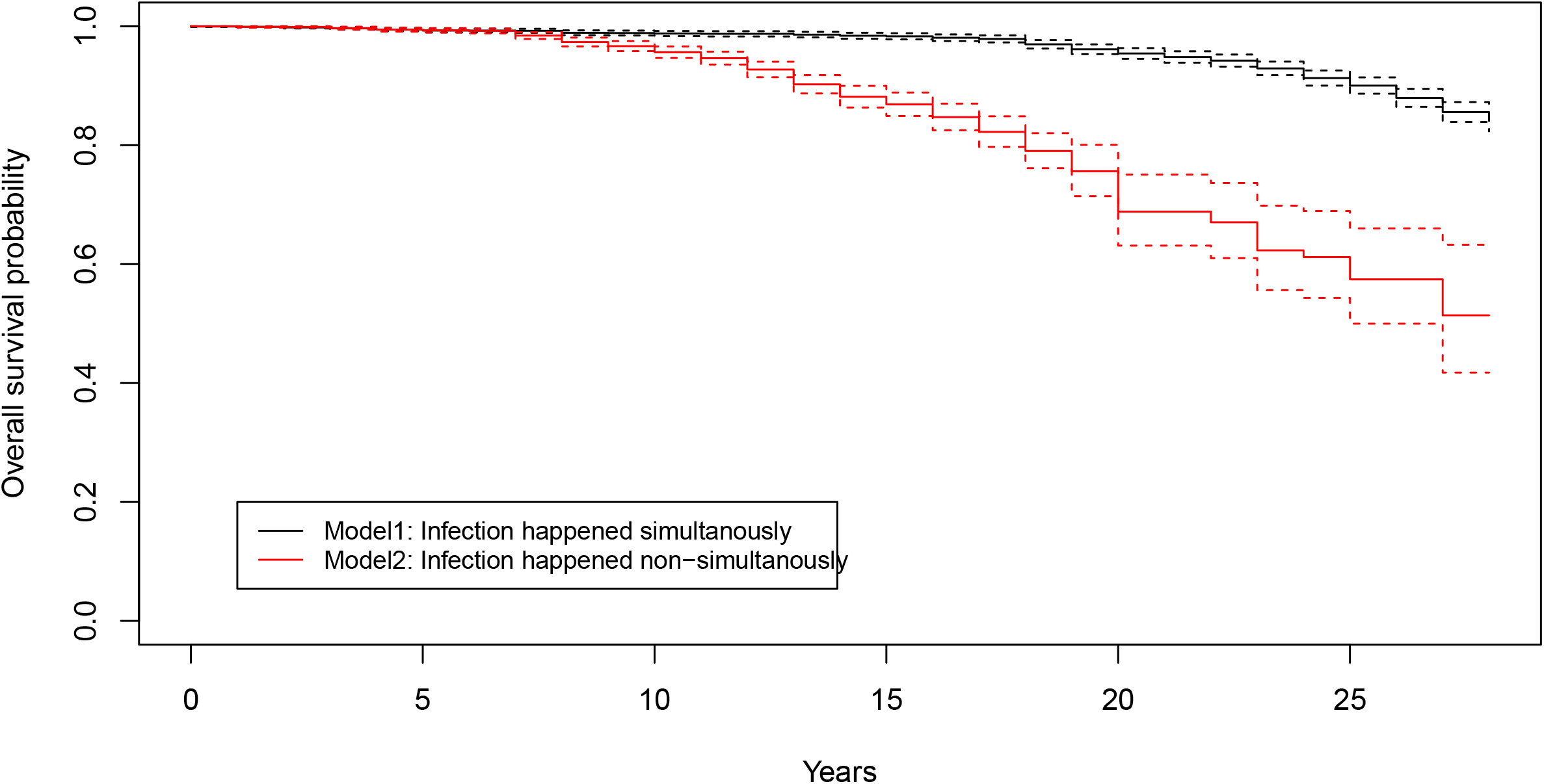
Kaplan-Maer survival function and overall survival probabilities between the two applied models. For Model 2 we assumed infection start of single countries according to dates of first arrival published in Vasaitis & Enderle 2017.

### 3.3. Cox proportional hazard model & Aalen additive regression model

Both the Cox-model and Aalens additive regression model unraveled significant covariates which determined survival of ash trees. The Cox-model found a significant influence of water status (p<0.01) and a highly significant influence of extreme precipitation events (p<0.001) as determinants for survival. Hazard ratios indicated that mortality risk increases towards sites having higher amount of excess water and also towards locations that have experienced more months with extreme precipitation during the last decades (Tab. 2). In concordance, the Aalen additive regression model revealed a strong negative impact of extreme precipitation events (slope: 0.003; p<0.01), but also a compensating effect of extreme drought events (slope: −0.012; p<0.01). Effect sizes of these two covariates were moderate and appeared mainly at time periods between 15-20 years after infection (Supporting information S1).

**Table 2:**
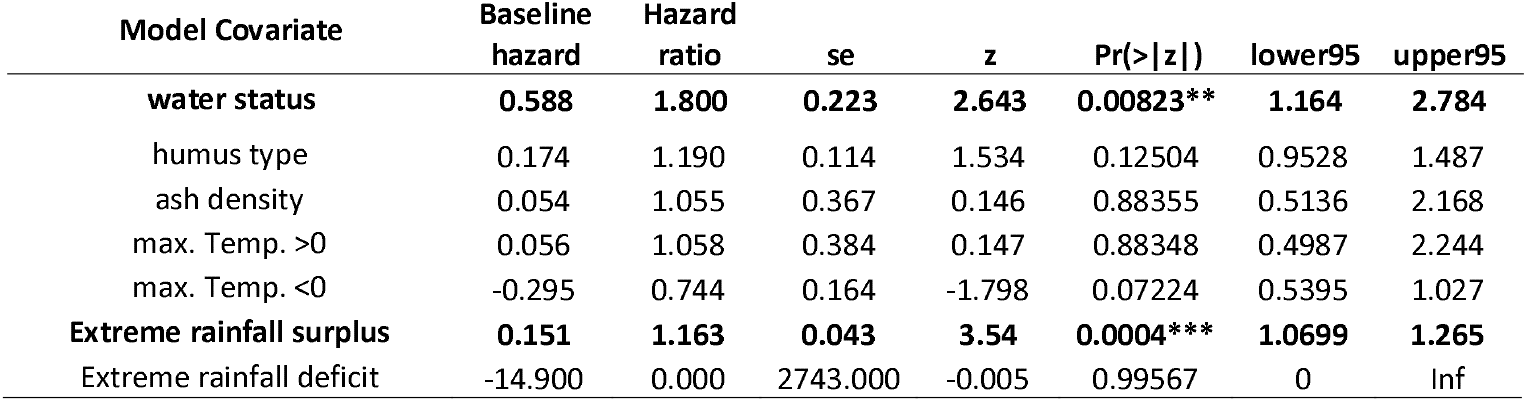
Summary statistics from the Cox hazrd model. Significant covariates are marked in bold.

| Model Covariate | Baseline hazard | Hazard ratio | se | z | Pr(>\|z\|) | lower95 | upper95 |
| --- | --- | --- | --- | --- | --- | --- | --- |
| water status | **0.588** | **1.800** | **0.223** | **2.643** | **0.00823\*\*** | **1.164** | **2.784** |
| humus type | 0.174 | 1.190 | 0.114 | 1.534 | 0.12504 | 0.9528 | 1.487 |
| ash density | 0.054 | 1.055 | 0.367 | 0.146 | 0.88355 | 0.5136 | 2.168 |
| max. Temp. >0 | 0.056 | 1.058 | 0.384 | 0.147 | 0.88348 | 0.4987 | 2.244 |
| max. Temp. <0 | -0.295 | 0.744 | 0.164 | -1.798 | 0.07224 | 0.5395 | 1.027 |
| **Extreme rainfall surplus** | **0.151** | **1.163** | **0.043** | **3.54** | **0.0004\*\*\*** | **1.0699** | **1.265** |
| Extreme rainfall deficit | -14.900 | 0.000 | 2743.000 | -0.005 | 0.99567 | 0 | Inf |

**Table 3:** Summary statistics from Aalens additive regression model. Significant covariates are marked in bold.

| **Model Covariate** | **slope** | **hazard** | **se** | **z** | **p** |
| --- | --- | --- | --- | --- | --- |
| Intercept | -0.024 | -0.002 | 0.001 | -1.895 | 0.058 |
| water status | 0.006 | 0.001 | 0.000 | 1.231 | 0.218 |
| humus type | 0.002 | 0.000 | 0.000 | 1.358 | 0.175 |
| ash density | -0.001 | 0.000 | 0.001 | 0.066 | 0.948 |
| **max. Temp. >0** | **-0.007** | **-0.001** | **0.000** | **-2.372** | **0.018** |
| max. Temp. <0 | -0.001 | 0.000 | 0.000 | 0.164 | 0.870 |
| **Extreme rainfall surplus** | **0.002** | **0.000** | **0.000** | **2.900** | **0.004** |
| **Extreme rainfall deficit** | **-0.014** | **-0.002** | **0.000** | **-4.278** | **0.000** |

### 3.4 Timer-series forecasting of crown condition development

Both estimators (ARIMA, exponential smoothing) gave similar forecasting results with continously increasing defoliation until the end of the forecasting period (2071) and strongly suggested that approximately until the year 2030 (exponential smoothing: 2031, ARIMA: 2033) mean defoliation could reach a critical threshold of 50% (Fig. 5). Confidence intervals were generally narrower for the ARIMA model compared to exponential smooting until 2036, but continously inreased after that time point towards more uncertain estimates. Both models were evaluated against the last ten years of observations (2010-2020) and exhibited high correlation between observed and forecasted data (Pearson-moment correlation 0.82 and 0.84, respectively). The ARIMA model showed generally better goodness-of-fit between observed and forecasted data with a MSE of 6.6% versus 16% (Supporting Information S2).

**Fig. 5:**
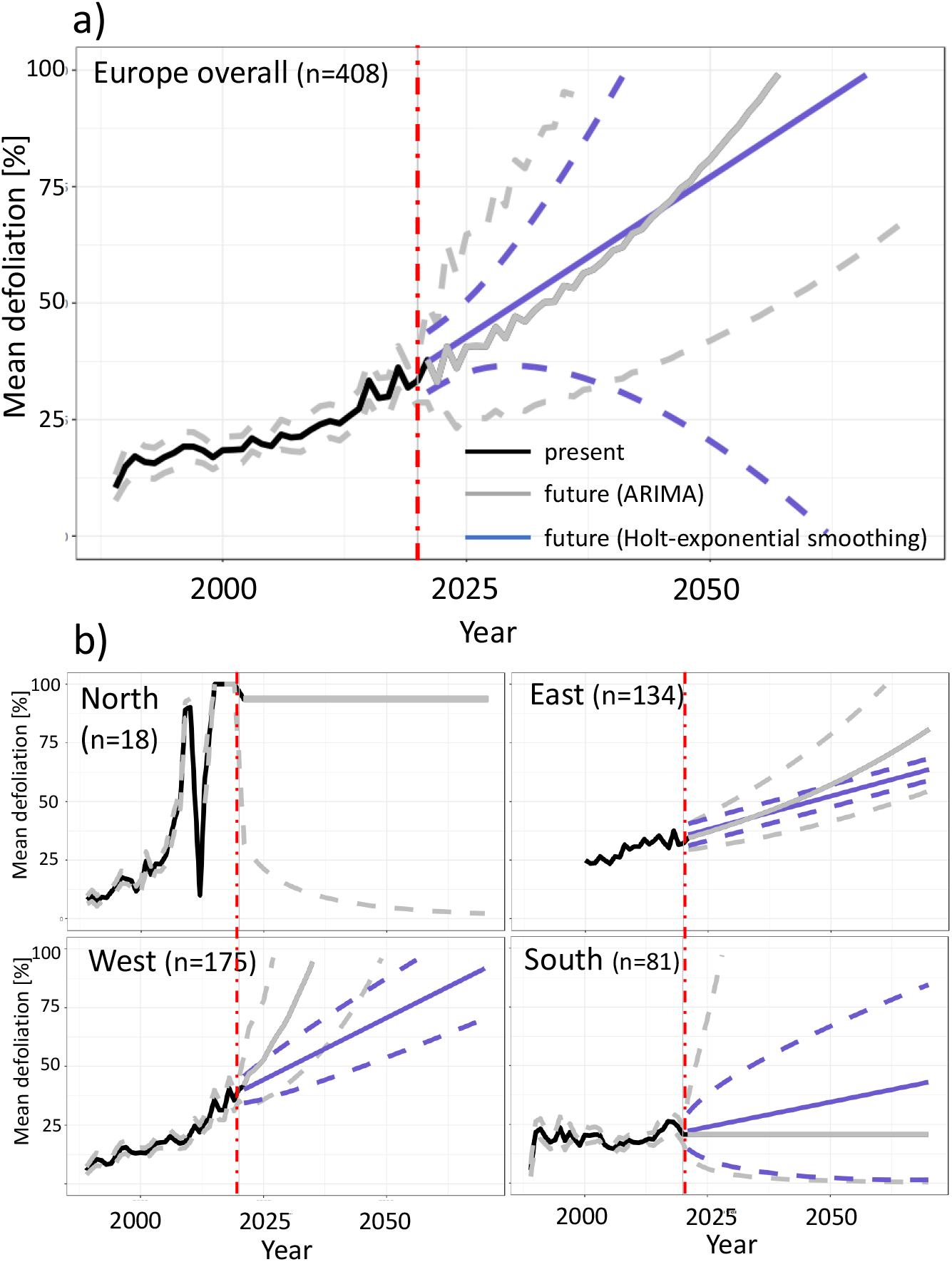
Time-series forecasting of mean defoliation per plot for a) entire Europe and b) at regional scale. Dashed lines show 95% confidence intervals.

### 4.4 Mortality of ash versus background mortality

Mortality of ash trees were largely decoupled from background mortality in the same plots (Fig. 6). As already outlined in section 2.2.1 some mortality occurred in southern Europe in 1990 probably as a result of abiotic factors and therefore strongly correlated with background mortality. Throughout the remaining period there was literally no correlation between mortality of ash and background mortality (r=0.15). Since 2010, ash mortality clearly accelerated and exceeded background mortality. By the end of 2020 10.8% of all surveyed ash trees had died while background mortality in the same plots was only 0.7%.

## 4. Discussion

### 4.1 Development in the last three decades and current hotspots of mortality

Mortality of both ash species has clearly accelerated during the last three decades and has already reached a catastrophic peak in northern Europe where almost all ash trees have disappeared from the survey plots by the end of 2020. This is clearly mirrored in the overall survival probability of 0.0 after nearly twenty years of infection history in this region. However, the rapid decrease of defoliation in Denmark in 2011 was simply caused by the fact that one survey plot harboured 22 ash trees in 2010 and all but one had 95% defoliation. Most likely the trees died or were cut in 2011 (but not reported dead to ICP Forests), and this plot was not longer surveyed in subsequent years so that literally only one ash tree in the last remaining danish plot existed by 2011. Our observations are in concordance with local and regional surveys from national forest inventories, which reported massive decline of European ash in Southern Scandinavia in recent years. Solheim & Hietala (2017) monitored the progress of ash dieback in south-western Norway since 2008 and estimated a spread velocity of 51 km per year. Diaz-Yanez *et al.* (2020) investigated survival of European ash in Norwegian forest inventory plots and found a 74% increase in mortality rate after the disease has become prevalent in Norway. Although the Norwegian plots in our study are generally charaterized by very low frequency of ash, the results can nevertheless be interpreted as a strong argument for the efficient and devastating spread of ash dieback, because the fungus seems not to depend on high host density in order to conquer new territory quickly. Also in Denmark and Sweden European ash began to disappear rapidly after the disease arrived in 2003 and 2001, respectively (Kjaer et al., 2017; Cleary et al., 2017). In Sweden *F. excelsior* became red-listed in 2010 and later even reached the status „critically-endangered” (Cleary et al. 2017). Consequently and in concert with the results presented here, the evidence strongly suggests that *Fraxinus excelsior* is currently under extreme extinction risk in northern Europe.

Furthermore, other hotspots of ash mortality are currently situated in the Baltic region (Lithuania, Poland), in eastern Europe (Belarus), north-eastern and south western Germany, as well as in eastern France (Fig. 2). Although not as severe as in in the northern part, our results clearly demonstrate that the disease is establishing across all of Europe, which was also very recently confirmed by national forest inventory data (e.g. Klesse et al. 2021). Since the Baltic and eastern countries were the first which came in contact with the disease in the beginning of the 1990s, the pattern may be unanticipated and one would expect highest mortality in these countries rather than in the north or in the west. One explanation for this could be that the density of survey plots in which ash occurrs are lower in the eastern part compared to the west (134 vs. 175). This is indeed true for Estonia and Latvia, for instance, where European ash is most likely underrepresented when compared to data from national forest inventories. The second explanation could be that assesment schemes in some eastern countries were harmonized a little later and therefore some trees were probably already gone when the survey started. As an example, Poland started to assess European ash first in 2006 when the disease had already been affecting stands for more than 15 years which could be the reason for the observed discrepancy.

Southern Europe seems to suffer least from ash dieback since mortality rates are still low and happened only sporadically. Mortality in northern Spain and southern France which occurred early at the beginning of the survey was certainly not related to ash dieback, since both areas are yet free of the disease (e.g. Trapiello et al. 2014). Besides the fact that the infection history of Southern Europe is still short and therefore may have caused higher survival, abiotic reasons such as higher temperatures, lower rainfall and generally drier site condtions could have contributed as well as they may inhibit fungal growth (e.g. Chumanowa et al. 2019; Grosdidier et al. 2018).

In general, mortality of ash during the last three decades was largely decoupled from background mortality and therefore unlikely to be caused by other biotic or abiotic reasons such as drought, storms, and others (Fig. 6). As already mentioned, only the very first incidences of mortality in 1990 were co-occurring with mortality in other species and therefore not attributable to ash dieback. Since 2010 the European-wide mortality of ash has even exceeded background mortality and is continously rising, while background mortality in the same plots remains constant. This demonstrates that in fact the majority of all cases in which ash trees are currently dying can be assigned to ash dieback as the predominant death cause. Although the specific crown damage cause in the ICP Forests Level I data can generally be retrieved down to the species level in the case of fungal agents (e.g. *Hymenoscyphus fraxineus* as the main causal agent of ash dieback), it is difficult to incorporate such data in detailed analyses, since the specific cause was hardly assessed and often not recognized in the very early years of the infection history and given that the damage coding system was first introduced in 2005. Nevertheless, *Hymenoscyphus fraxineus* (or *Chalara fraxinea* as its asexual stadium is named) is currently accounting for 53% of all damage causes where a specific biotic agent was determined (data not shown). This corresponds to approximately every fourth ash tree in the survey (23.4%) which is already carrying the disease by 2020.

**Fig. 6:**
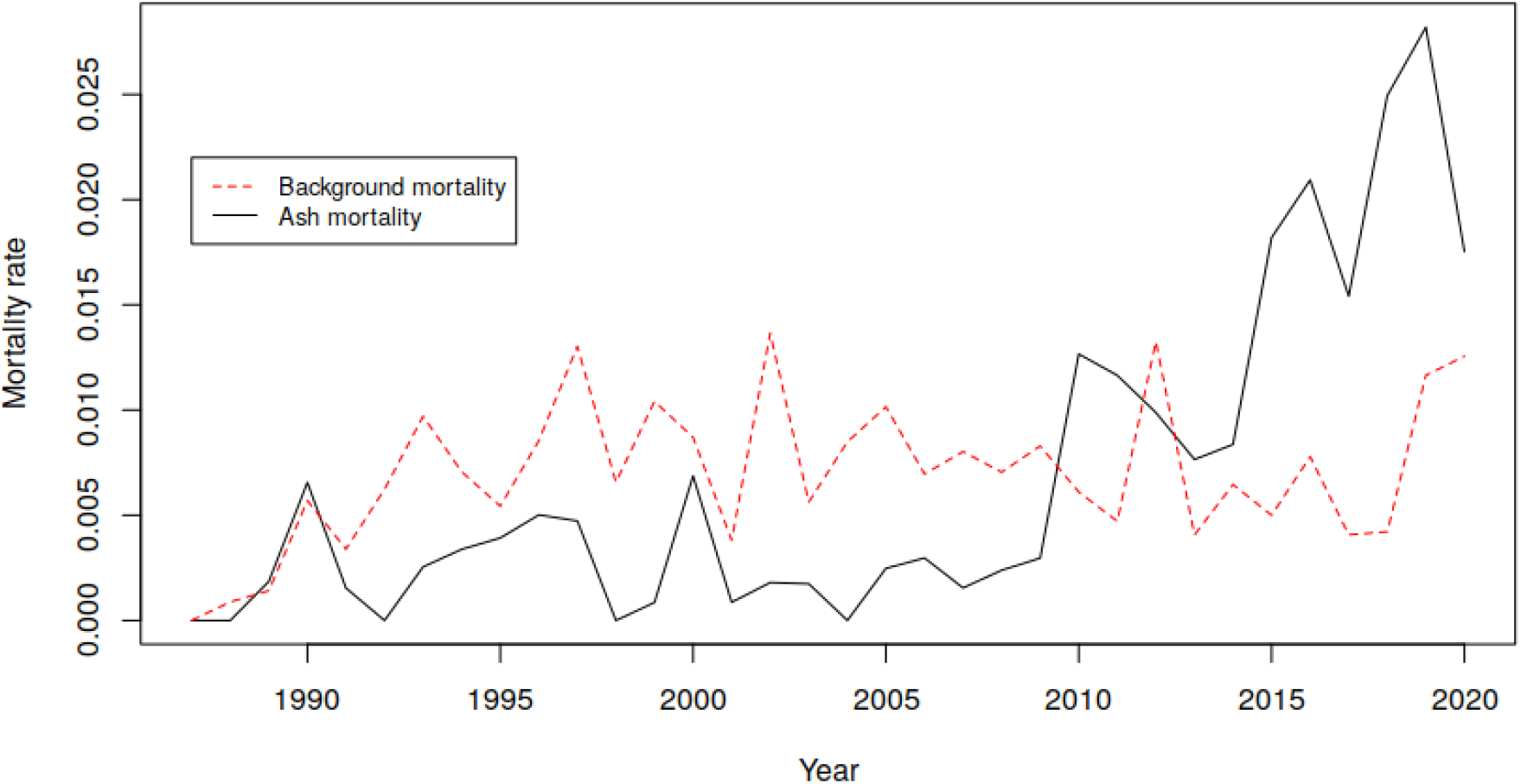
Correlation between ash mortality and background mortality by survey year. Note that data was aggregated over all plots regardless of country or region for purposes of illustration.

### 4.2 Overall survival probability after three decades of ash dieback exposure

We aimed to assess the overall survival probability of ash after an exposure time of 28 years to ash dieback in order to provide meaningful planning horizons for conservation and ecosystem management. We found that the probability of survival was much lower when the exposure time - approximated by the first observation of ADB at national levels - was corrected for the infection history, since the disease spread concentrically from north-eastern Poland to other regions of Europe. The assumption that entire Europe was simultanously infected in 1992 is too simplistic and would have caused much longer time spans from infection to death for regions further apart from the epicentre. This, in turn, generated much higher survival probabilities compared to the more realistic scenario of shifted infection history.

We estimated that the overall survival probability of ash after nearly three decades of exposure is approximately 50%, but with large differences among regions (e.g. northern group: 0.0, southern group: 0.9). Coker et al. (2019) estimated that mortality of ash in woodlands after 11 years of exposure reaches a maximum at 60% and remains constant afterwards. Although the values from Coker et al. (2019) and those from our study cannot directly be compared due to methodological differences, such a trend cannot be derived from our results. In contrast, the survival probability curve in our study shows a constant downward trend even after 27 years of exposure and the evidence strongly suggests that it will be continued until host availability has reached a value critical for the survival of the pathogen.

### 4.3 Environmental drivers of survival

We found significant site parameters critical for survival that were also unraveled in earlier studies such as water status. Trees growing in locations with moister growing conditions are under higher risk to die compared to those growing in drier locations. The hazard ratio of 1.8 suggests that, for example, trees growing at sites with excessive water status have a 1.8-fold higher risk of death compared to trees growing under sufficient water status and a 3.6-fold higher risk compared to trees growing under insufficient or dry conditions. In accordance, trees growing at sites which experienced one more months with extreme rainfall surplus during the observation period had a 1.07 fold higher risk. Both covariates are likely to favour the disease, since moisture is a critical factor during infection and for the efficient spread of ascospores (Enderle et al. 2019). Excess moisture may also plays a critical role for growth of other wood-deacaying fungi such as *Armillaria spec,* which were recently found to be involved in subsequent root rot of ash trees after infection with ADB (e.g. Chandelier et al. 2016).

Aalen’s additive regression model also unraveled months with extreme high temperature as well as months with extreme rainfall deficit as factors which reduce the risk of death. While this pattern is largely corroborated by other studies (e.g. Grosdidier et al. 2018), the effect sizes of both covariates in our model were only moderate and hence it desires more research in order to confirm whether this relationship holds true at continental scale.

Lower host abundance was recently associated with lower mortality in Swiss ash stands (Klesse et al. 2021), but our study did not confirm this relationship. At continental scale ash trees were killed regardless of whether they occurred at high abundance or low abundance. This demonstrates once more the devastating and highly efficient spreading capacity of the pathogen which allows the fungus to enter new territory at a velocity of 50-70 km per year (Gross et al. 2014; Solheim & Hietala 2017).

### 4.5. Appeal for a joint conservation strategy at European level

Based on the ICP Forests Level I crown defoliation survey we show the value of such data for monitoring ash dieback at continental scale. Although *F. excelsior* and *F. angustifolia* are minor tree species in most of the survey countries, they nevertheless constitute keystone species with high importance for ecological communities and local economy (Enderle et al. 2018; Mitchell et al., 2014). In the British Isles and Ireland, the importance of ash is even higher as *F. excelsior* is occupying a larger proportion of woodlands compared to other countries (Clark & Webber, 2017, Dandy et al. 2017). Unfortunately, the United Kingdom never assessed ash for defoliation and left the ICP Forests Level I network in 2011 so that no data on ash dieback is available from there.

Overall our study indicates a strongly progressive pattern of ash decline across the last three decades and we estimated that an Europe-wide average defoliation of 50% could be reached as early as 2030. However, we also outlined that the disease strongly differs among regions, since the progress seems to be slower in the south compared to the north. The threshold of 50% defoliation can be seen indeed as a critical benchmark, since we estimated that trees that have reached this defoliation status died within the following 9 years (data not shown). A further decline without joint rescue efforts at European level could cause non-reversible loss of genetic diversity which in turn could cause a critical minimum population size vital for long-term survival. Although a minor proportion of ash trees is likely inheriting resistance against ADB (Plumb et al. 2019; Kjær et al., 2012), this alone will not guarantee that the ash population will recover after the disease has reached an equilibrium state. Evans (2019) simulated that long-term recovery of ash is highly dependent on the proportion of ash trees carrying natural resistance and secondly on the degree of heritability of resistance. However, even under extreme high heritability assumptions, long-term recovery under natural conditions will remain low when the founder population consists of only few trees. Hence, without efforts at European level it is very likely that the two ash species *F. excelsior* and *F. angutifolia* will face the same fate as dutch elm, which largely disappeared in most parts of Europe after the devastating dutch elm disease (Buggs, 2020; Tomlinson & Potter, 2010).

Conservation efforts at national level have recently been undertaken in Great Britain, Ireland, Denmark, and Austria and new projects are currently starting in Germany and other countries. For instance, reference genomes for *F. excelsior* and *F. angustifolia* have been completely sequenced (e.g. Sollars et al. 2017 and http://www.ashgenome.org/, 2021) and novel markers which are able to discriminate between resistent and susceptible *F. excelsior* trees have been developed (Stocks et al. 2019). On the other hand, progeny tests which aim to evaluate resistance against ADB of several thousand saplings of European ash harvested from field-resistant mother trees are currently being tested in Austria (http://www.esche-in-not.at/index.php). Monitoring ADB will become crucial in the future in order to see whether undertaken efforts have been successful, but also to identify regions where immediate actions are required as has been demonstarted in this study. In this regard, remotely sensed resources at high temporal and spatial resolution need to be developed and evaluated and can be combined with ground-truthed survey data such as the ICP Forests Level I dataset. We strongly believe that when applied in a Pan-European project consortium all these extremely valuable and already gathered datasets (molecular genetics and genomic markers, quantitative breeding & progeny testing, monitoring & remote sensing, etc.) can significantly contribute in order to avoid that ash will largely disappear from the European forest landscape.

## 6. Acknowledgements

This study is carried out with the support of the European Regional Development Fund and the programme Mobilitas PLUSS (Project ID: MOBJD588) granted by the Estonian Research Agency (ETAG). The evaluation was based on data collected by partners of the official UNECE ICP Forests Network (http://icp-forests.net/contributors). Part of the data was co-financed by the European Commission (Data achieved at 12.04.2021).

## References

Aalen, O. O. (1989). A linear regression model for the analysis of life times. Statistics in medicine, 8(8), 907–925.

Børja, I., Timmermann, V., Hietala, A. M., Tollefsrud, M. M., Nagy, N. E., Vivian-Smith, A.,… & Solheim, H. (2017). Ash dieback in Norway–current situation. Dieback of European Ash (Fraxinus spp.):Consequences and Guidelines for Sustainable Management, 166–175.

Buggs RJA. Changing perceptions of tree resistance research. Plants, People, Planet. 2020;2:2–4. https://doi.org/10.1002/ppp3.10089

Chandelier A, Gerarts F, San Martin G, Herman M, Delahaye L. Temporal evolution of collar lesions associated with ash dieback and the occurrence of Armillaria in Belgian forests. Forest Pathology 2016;46(4):289–97. doi: 10.1111/efp.12258.

Chumanová, E., Romportl, D., Havrdová, L., Zahradník, D., Pešková, V., & Černý, K. (2019). Predicting ash dieback severity and environmental suitability for the disease in forest stands. Scandinavian Journal of Forest Research, 34(4), 254–266.

Clark J, Webber J. The ash resource and the response to ash dieback in Great Britain. In: Vasaitis R, Enderle R, editors. Dieback of European ash (Fraxinus spp.) - Consequences and Guidelines for Sustainable Management. Uppsala, Sweden: Swedish University of Agricultural Sciences; 2017. p. 228–37.

Cleary, M., Nguyen, D., Stener, L. G., Stenlid, J., & Skovsgaard, J. P. (2017). Ash and ash dieback in Sweden: A review of disease history, current status, pathogen and host dynamics, host tolerance and management options in forests and landscapes. Dieback of European Ash (Fraxinus spp.}: Consequences and Guidelines for Sustainable Management, 195–208.

Coker, T. L., Rozsypálek, J., Edwards, A., Harwood, T. P., Butfoy, L., & Buggs, R. J. (2019). Estimating mortality rates of European ash (Fraxinus excelsior) under the ash dieback (Hymenoscyphus fraxineus) epidemic. Plants, People, Planet, 1(1), 48–58.

Cox, D. R. (1972). Regression models and life-tables. Journal of the Royal Statistical Society: Series B (Methodological), 34(2), 187–202.

Dandy N, Marzano M, Porth EF, Urquhart J, Potter C. Who has a stake in ash dieback? A conceptual framework for the identification and categorisation of tree health stakeholders. In: Vasaitis R, Enderle R, editors. Dieback of European ash (Fraxinus spp.) – Consequences and Guidelines for Sustainable Management. Uppsala, Sweden: Swedish University of Agricultural Sciences; 2017. p. 15–26.

Díaz-Yáñez, O., Mola-Yudego, B., Timmermann, V., Tollefsrud, M. M., Hietala, A. M., & Oliva, J. (2020). The invasive forest pathogen Hymenoscyphus fraxineus boosts mortality and triggers niche replacement of European ash (Fraxinus excelsior). Scientific Reports, 10(1), 1–10.

Dufour, S., & Piégay, H. (2008). Geomorphological controls of Fraxinus excelsior growth and regeneration in floodplain forests. Ecology, 89(1), 205–215.

Eichhorn J, Roskams P, Potočić N, Timmermann V, Ferretti M, Mues V, Szepesi A, Durrant D, Seletković I, Schröck H-W, Nevalainen S, Bussotti F, Garcia P, Wulff S, 2016: Part IV: Visual Assessment of Crown Condition and Damaging Agents. In: UNECE ICP Forests Programme Co-ordinating Centre (ed.): Manual on methods and criteria for harmonized sampling, assessment, monitoring and analysis of the effects of air pollution on forests. Thünen Institute of Forest Ecosystems, Eberswalde, Germany, 49 p. + Annex [http://www.icp-forests.org/manual.htm].

Enderle, R., Stenlid, J., & Vasaitis, R. (2019). An overview of ash (Fraxinus spp.) and the ash dieback disease in Europe. CAB Rev, 14, 1–12.

Enderle, R., Metzler, B., Riemer, U., & Kändler, G. (2018). Ash dieback on sample points of the national forest inventory in south-western Germany. Forests, 9(1), 25.

Evans, M. R. (2019). Will natural resistance result in populations of ash trees remaining in British woodlands after a century of ash dieback disease?. Royal Society Open Science, 6(8), 190908.

Ghelardini, L., Migliorini, D., Santini, A., Pepori, A. L., Maresi, G., Vai, N.,… & Luchi, N. (2017). From the Alps to the Apennines: possible spread of ash dieback in Mediterranean areas. Dieback of European Ash (Fraxinus spp.): Consequences and Guidelines for Sustainable Management, 140–149.

Godaert, L, Bartholet, S., Dorléans, F., Najioullah, F., Colas, S., Fanon, J. L.,… & Dramé, M. (2018). Prognostic factors of inhospital death in elderly patients: a time-to-event analysis of a cohort study in Martinique (French West Indies). BMJ open, 8(1), e018838.

Grosdidier M, Ioos R, Marçais B. Do higher summer temperatures restrict the dissemination of Hymenoscyphus fraxineus in France? Forest Pathology 2018;48(4):e12426. doi: 10.1111/efp.12426

Gross, A., Holdenrieder, O., Pautasso, M., Queloz, V., & Sieber, T. N. (2014). H ymenoscyphus pseudoalbidus, the causal agent of E uropean ash dieback. Molecular Plant Pathology, 15(1), 5–21.

Haylock, M. R., Hofstra, N., Klein Tank, A. M. G., Klok, E. J., Jones, P. D., & New, M. (2008). A European daily high-resolution gridded data set of surface temperature and precipitation for 1950-2006. Journal of Geophysical Research: Atmospheres, 113(D20).

Heinze, B., Tiefenbacher, H., Litschauer, R., & Kirisits, T. (2017). Ash dieback in Austria—history, current situation and outlook. Dieback of European Ash (Fraxinus spp.)–Consequences and Guidelines for Sustainable Management, 33–52.

Hill, L., Jones, G., Atkinson, N., Hector, A., Hemery, G., & Brown, N. (2019). The£ 15 billion cost of ash dieback in Britain. Current Biology, 29(9), R315–R316.

Holt, C.C. (1957). Forecasting trends and season-als by exponentially weighted averages, Carnegie Institute of Technology, Pittsburgh ONR memorandum no. 52

Hyndman, R. J., & Khandakar, Y. (2008). Automatic time series forecasting: the forecast package for R. Journal of statistical software, 27(3), 1–22.

Kjær ED, McKinney LV, Hansen LN, Olrik DC, Lobo A, Thomsen IM, et al. Genetics of ash dieback resistance in a restoration context – experiences from Denmark. In: Vasaitis R, Enderle R, editors. Dieback of European ash (Fraxinus spp.) – Consequences and Guidelines for Sustainable Management. Uppsala, Sweden: Swedish University of Agricultural Sciences; 2017. p. 106–14.

Kjær, E. D., McKinney, L. V., Nielsen, L. R., Hansen, L. N., & Hansen, J. K. (2012). Adaptive potential of ash (Fraxinus excelsior) populations against the novel emerging pathogen Hymenoscyphus pseudoalbidus. Evolutionary Applications, 5(3), 219–228.

Klesse, S., Abegg, M., Hopf, S. E., Gossner, M. M., Rigling, A., & Queloz, V. (2021). Spread and severity of ash dieback in Switzerland–tree characteristics and landscape features explain varying mortality probability. Frontiers in Forests and Global Change, 4, 18.

Klesse, S., von Arx, G., Gossner, M. M., Hug, C., Rigling, A., & Queloz, V. (2020). Amplifying feedback loop between growth and wood anatomical characteristics of Fraxinus excelsior explains size-related susceptibility to ash dieback. Tree Physiology, 41(5), 683–696.

Koontz, M. J., Latimer, A. M., Mortenson, L. A., Fettig, C. J., & North, M. P. (2021). Cross-scale interaction of host tree size and climatic water deficit governs bark beetle-induced tree mortality. Nature Communications, 12(1), 1–13.

Marçais, B., Husson, C., Godart, L., & Cael, O. (2016). Influence of site and stand factors on Hymenoscyphus fraxineus-induced basal lesions. Plant Pathology, 65(9), 1452–1461.

Mitchell, R. J., Beaton, J. K., Bellamy, P. E., Broome, A., Chetcuti, J., Eaton, S.,… & Woodward, S. (2014). Ash dieback in the UK: a review of the ecological and conservation implications and potential management options. Biological conservation, 175, 95–109.

Neumann, M., Mues, V., Moreno, A., Hasenauer, H., & Seidl, R. (2017). Climate variability drives recent tree mortality in Europe. Global Change Biology, 23(11), 4788–4797.

Orton, E. S., Brasier, C. M., Bilham, L. J., Bansal, A., Webber, J. F., & Brown, J. K. (2018). Population structure of the ash dieback pathogen, Hymenoscyphus fraxineus, in relation to its mode of arrival in the UK. Plant pathology, 67(2), 255–264.

Pliûra, A., & Heuertz, M. (2003). EUFORGEN Technical Guidelines for genetic conservation and use for common ash (Fraxinus excelsior). Bioversity International.

Plumb, W. J., Coker, T. L., Stocks, J. J., Woodcock, P., Quine, C. P., Nemesio-Gorriz, M.,… & Buggs, R. J. (2020). The viability of a breeding programme for ash in the British Isles in the face of ash dieback. Plants, People, Planet, 2(1), 29–40.

Przybył, K. (2002). Fungi associated with necrotic apical parts of Fraxinus excelsior shoots. Forest Pathology, 32(6), 387–394.

Queloz, V., Hopf, S., Schoebel, C. N., Rigling, D., & Gross, A. (2017). Ash dieback in Switzerland: history and scientific achievements. Dieback of European ash, 68–78.

R Development Core Team. (2017). RStudio, R: A Language and Environment for Statistical Computing.

Sargeran, K., Murtomaa, H., Safavi, S. M. R., Vehkalahti, M. M., & Teronen, O. (2008). Survival after diagnosis of cancer of the oral cavity. British Journal of Oral and Maxillofacial Surgery, 46(3), 187–191.

Senf, C., Buras, A., Zang, C. S., Rammig, A., & Seidl, R. (2020). Excess forest mortality is consistently linked to drought across Europe. Nature Communications, 11(1), 1–8.

Solheim, H. & Hietala, AM. (2017). Spread of ash dieback in Norway. Baltic Forestry 23(1):1–6

Sollars, E. S., Harper, A. L., Kelly, L. J., Sambles, C. M., Ramirez-Gonzalez, R. H., Swarbreck, D.,… & Buggs, R. J. (2017). Genome sequence and genetic diversity of European ash trees. Nature, 541(7636), 212–216.

Stocks, J. J., Metheringham, C. L., Plumb, W. J., Lee, S. J., Kelly, L. J., Nichols, R. A., & Buggs, R. J. (2019). Genomic basis of European ash tree resistance to ash dieback fungus. Nature ecology & evolution, 3(12), 1686–1696.

Stocks, J. J., Buggs, R. J., & Lee, S. J. (2017). A first assessment of Fraxinus excelsior (common ash) susceptibility to Hymenoscyphus fraxineus (ash dieback) throughout the British Isles. Scientific Reports, 7(1), 1–7.

Taccoen, A., Piedallu, C., Seynave, I., Gégout-Petit, A., Nageleisen, L. M., Bréda, N., & Gégout, J. C. (2021). Climate change impact on tree mortality differs with tree social status. Forest Ecology and Management, 489, 119048.

Therneau, T. (2020). A package for survival analysis in R. Available under: [URL: https://cran.r-project.org/web/packages/survival/vignettes/survival.pdf] last request: May 26th 2021

Therneau, T. M., & Grambsch, P. M. (2000). The cox model. In Modeling survival data: extending the Cox model (pp. 39–77). Springer, New York, NY.

Timmermann, V., Potočić, N., Ognjenović, M. & Kirchner, T. (2021). Tree crown condition in 2020. In: Michel, A., Prescher, A.K. & Schwärzel, K. (eds.) 2021. Forest Condition in Europe: The 2021 Assessment. ICP Forests Technical Report under the UNECE Convention on Long-range Transboundary Air Pollution (Air Convention). Eberswalde: Thünen Institute. In prep.

Tomlinson, I., & Potter, C. (2010). ‘Too little, too late’? Science, policy and Dutch Elm Disease in the UK. Journal of Historical Geography, 36(2), 121–131.

Trapiello, E., Schoebel, C. N., & Rigling, D. (2017). Fungal community in symptomatic ash leaves in Spain. Baltic Forestry, 23(1), 68–73.

Vasaitis, R., & Enderle, R. (2017). Dieback of European ash (Fraxinus spp.) – consequences and guidelines for sustainable management. Dieback of European ash (Fraxinus spp.)-consequences and guidelines for sustainable management. Report on COST Action FP1103 FRAXBACK. ISBN 978-91-576-8696-1. SLU Swedish University of Agricultural Sciences.

