## Supplementary file S1 for "European-wide forest monitoring substantiate the neccessity for a joint conservation strategy to rescue European ash species (*Fraxinus spp*.)"

### Slide 1
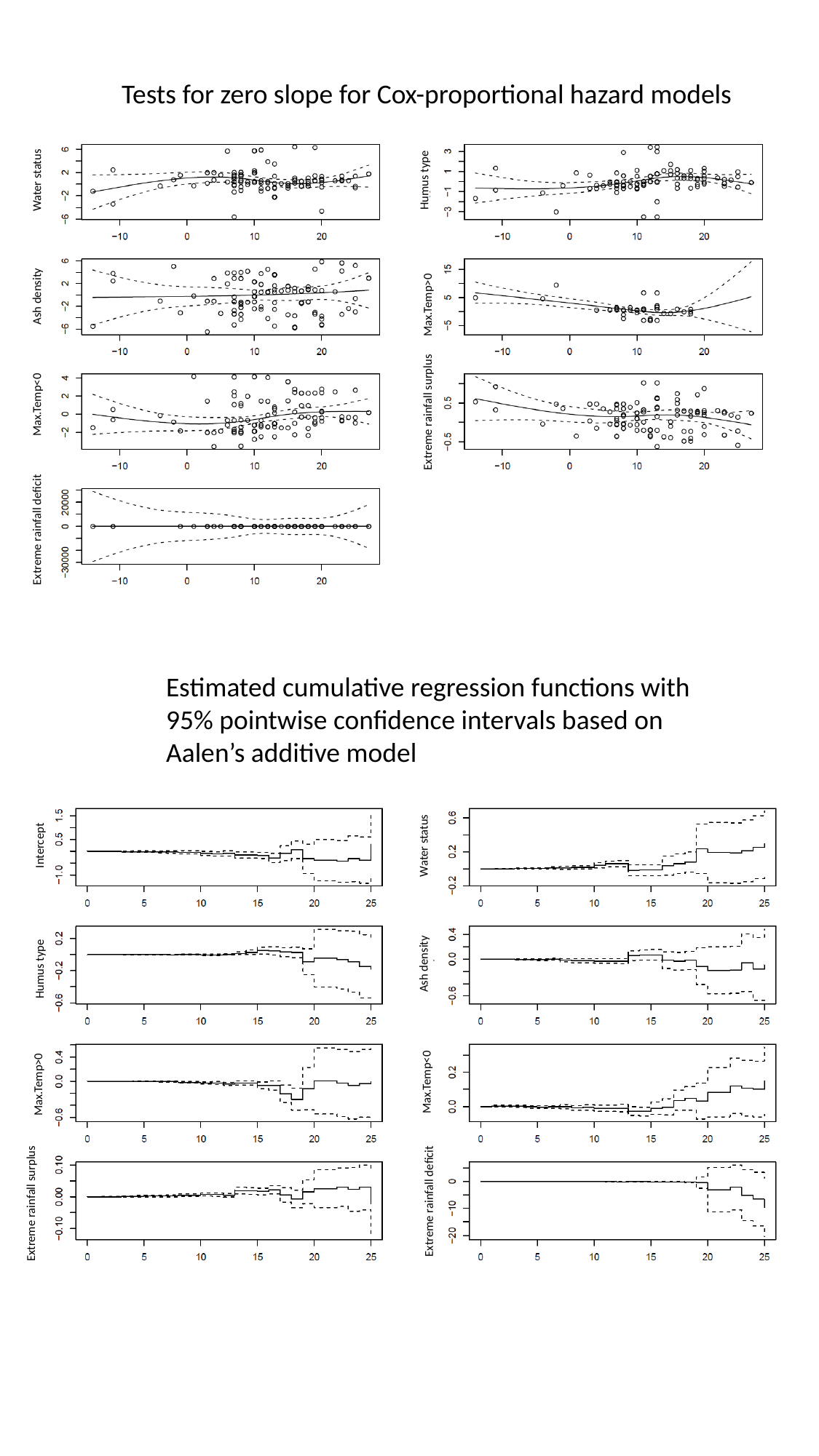

Tests for zero slope for Cox-proportional hazard models
Water status
Humus type
Ash density
Max.Temp>0
Max.Temp<0
Extreme rainfall surplus
Extreme rainfall deficit
Estimated cumulative regression functions with 95% pointwise confidence intervals based on Aalen’s additive model
Intercept
Water status
Ash density
Humus type
Max.Temp<0
Max.Temp>0
Extreme rainfall deficit
Extreme rainfall surplus
