## Supplementary file S2 for "European-wide forest monitoring substantiate the neccessity for a joint conservation strategy to rescue European ash species (*Fraxinus spp*.)"

### Slide 1
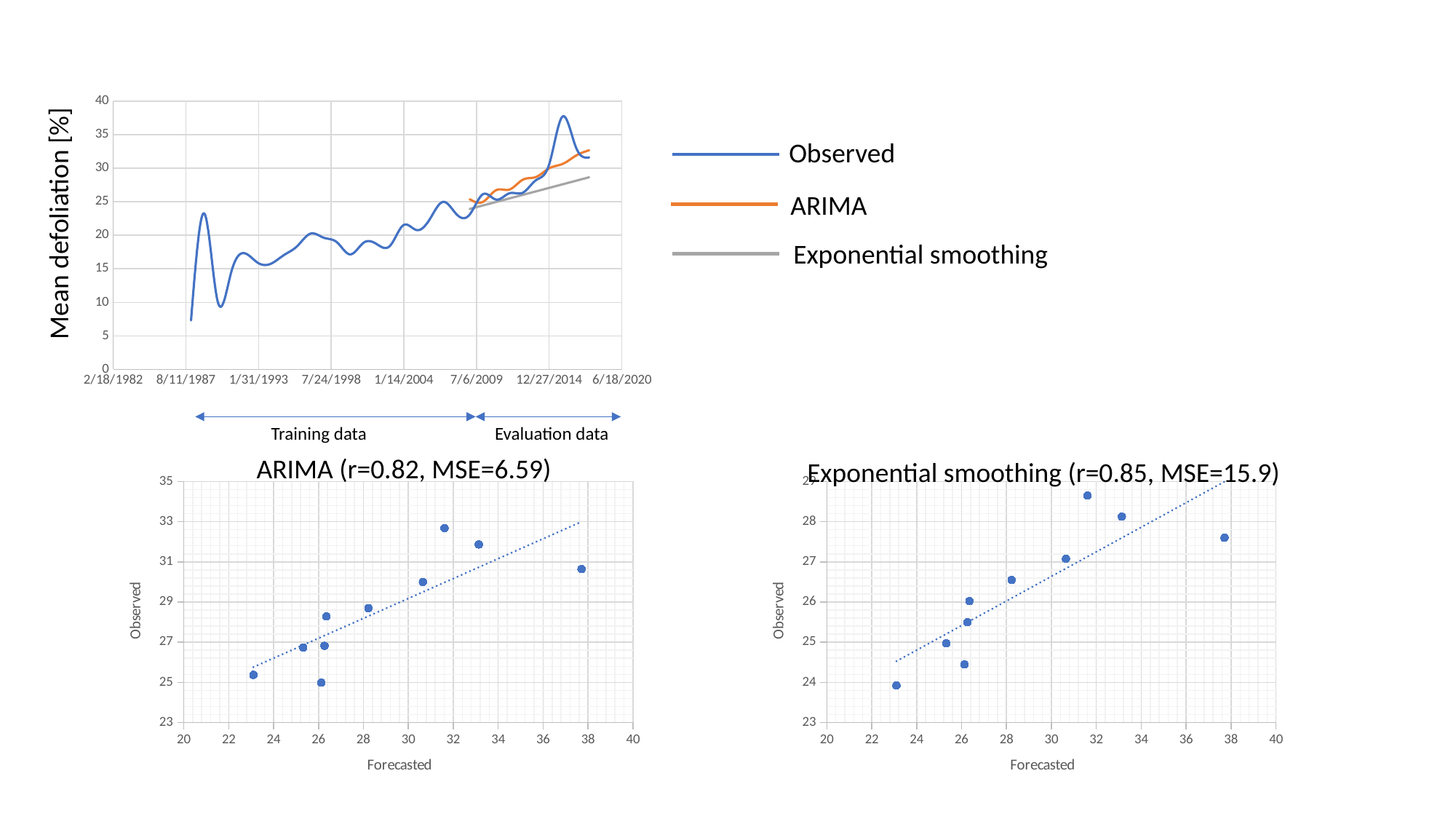

#### Chart
| Category | | | |
|---|---|---|---|Observed
ARIMA
Mean defoliation [%]
Exponential smoothing
Training data
Evaluation data
ARIMA (r=0.82, MSE=6.59)
Exponential smoothing (r=0.85, MSE=15.9)
#### Chart
| Category |
|---|
#### Chart
| Category |
|---|
